# Finding regulatory modules of chemical reaction systems

**DOI:** 10.1101/2023.10.12.561372

**Authors:** Yuhei Yamauchi, Atsuki Hishida, Takashi Okada, Atsushi Mochizuki

## Abstract

Within a cell, numerous chemical reactions form chemical reaction networks (CRNs), which are the origins of cellular functions. We previously developed a theoretical method called structural sensitivity analysis (SSA) [1], which enables us to determine, solely from the network structure, the qualitative changes in the steady-state concentrations of chemicals resulting from the perturbations to a parameter. Notably, if a subnetwork satisfies specific topological conditions, it is referred to as a buffering structure, and the effects of perturbations to the parameter within the subnetwork are localized to the subnetwork (the law of localization) [2, 3]. A buffering structure can be the origin of modularity in the regulation of cellular functions generated from CRNs. However, an efficient method to search for buffering structures in a large CRN has not yet been established. In this study, we proved the “inverse theorem” of the law of localization, which states that a certain subnetwork exhibiting a confined response range is always a buffering structure. In other words, we are able to identify buffering structures in terms of sensitivities rather than the topological conditions. By leveraging this property, we developed an algorithm to enumerate all buffering structures for a given network by calculating the sensitivity. In addition, using the inverse theorem, we demonstrated that the hierarchy among nonzero responses is equivalent to the hierarchy of buffering structures. Our method will be a powerful tool for understanding the regulation of cellular functions generated from CRNs.

## 1 Introduction

In living cells, chemical reactions are connected by sharing their products and substrates, forming chemical reaction networks (CRNs), such as metabolic networks and signal transduction networks. The dynamical behavior of chemicals derived from these CRNs underlies physiological functions, such as metabolism, cell cycle control, and signal transduction [4, 5]. Each reaction is regulated by enzyme activity, and changes in enzyme activity cause dynamical changes in the concentration of each chemical in the system. It is widely believed that cells regulate physiological functions through the modulation of enzyme activity [6, 7, 8, 9]. Various physiological functions can arise even from a single CRN. For instance, a metabolic network is composed of many pathways, such as carbon metabolism and the amino acid synthesis pathway [10, 11]. Since different pathways can share the same chemicals, these pathways are interconnected with each other, forming a single connected metabolic network [12, 13, 14]. It is unclear how different pathways are regulated separately because the modulation of one pathway can affect other pathways.

We previously developed a theoretical framework that may provide an answer to this question [2, 3]. This is based on structural sensitivity analysis (SSA) [1], which allows us to determine, solely from the structure of the CRN, the qualitative changes (no change, increase, or decrease) in steady-state concentrations of each chemical in response to the modulation of enzyme activity. In SSA, the parameters are either the activities of the enzymes catalyzing the reactions (reaction rate parameters) or the conserved quantities. The concept of “buffering structures” derived from SSA is important for understanding the range within which the effects of changes in parameters propagate within a CRN in terms of network topology [2, 3]. A subnetwork in a CRN, consisting of a subset of chemicals and a subset of reactions, is called a buffering structure when it satisfies the following two conditions: (i) The subnetwork contains all reactions whose reaction rates depend on the concentrations of chemicals within it (“output-complete”). (ii) The index, defined by the −(the number of chemicals)+(the number of reactions)−(the number of cycles)+(the number of conserved quantities), is equal to zero. It has been mathematically demonstrated that the steady-state responses to the perturbation of a parameter within a buffering structure are confined within the buffering structure. When distinct non-overlapping buffering structures (denoted by Γ_1_ and Γ_2_) coexist within a network, the perturbations to parameters within Γ_1_ do not affect the steady-state concentration of chemicals within Γ_2_, and vice versa. In this sense, independent regulation between distinct buffering structures is achieved. Taking biological networks for example, the tricarboxylic acid (TCA) cycle and the pentose phosphate pathway (PPP) in metabolic networks are contained in separate buffering structures [2], indicating that it is possible to modulate the TCA cycle without affecting the PPP pathway, and vice versa. A buffering structure represents a novel concept in CRNs: a “regulatory module” can naturally arise from topology of a network. Finding all the buffering structures in a given CRN will be important for clarifying how different functions arising from a CRN are regulated.

In [2], it is pointed out that the existence of buffering structures can provide the system with a property called “perfect adaptation”. A system is said to exhibit adaptation if it shows a transient response to a stimulus and goes back to the original state [15, 16, 17, 18]. A well-known example is the response of bacteria to changes in nutrient concentration during chemotaxis [19, 20, 21]. In particular, if, after a perturbation to a parameter, the system eventually returns to its original state exactly, it is called perfect adaptation, which has been studied theoretically [22].

While exploring the buffering structures in a CRN is important, an efficient method to search for all of them has not been established. One approach to search for buffering structures is to check, for all subnetworks of CRNs, if the aforementioned two conditions are satisfied (the “brute-force method”). However, as the size of the CRN increases, the number of candidate subnetworks grows significantly, leading to a substantial increase in the computational cost. In previous studies [2, 3, 23], an ad-hoc method was employed where buffering structures are identified by searching for candidate subnetworks that show confined responses, using the calculation of sensitivities. However, it remained unclear whether all subnetworks calculated from the method satisfy the topological conditions of buffering structures.

In this study, we proved the “inverse theorem” of the law of localization, which states that an output-complete subnetwork exhibiting a confined response range (“response block”) is always a buffering structure. This ensures that buffering structures can be exhaustively identified by all of the response blocks, which are obtained via the sensitivity calculation. The proposed method is much more efficient than the brute-force method. We implemented the algorithm as Python pipelines, called ibuffpy (https://github.com/hishidagit/SSApy).

We previously found that the nonzero response patterns under perturbations of different parameters can exhibit inclusion relations among them, i.e., they exhibit hierarchical structures [2]. In addition, buffering structures have been suggested to exhibit inclusion relations [2]. Based on the inverse theorem, we showed that a buffering structure and a confined response have a one-to-one correspondence. We thus understand that nonzero response patterns always show hierarchical patterns. Furthermore, we propose an algorithm to depict this hierarchy graphically through the computation of buffering structures, which is also implemented in ibuffpy.

The paper is organized as follows. In Section 2, we briefly review SSA and the concept of a buffering structure presented in [1, 2]. In Section 3, we prove our main theorem: the inverse theorem of the law of localization. In Section 4, we present an efficient algorithm to exhaustively find buffering structures. In Section 5, we show the equivalence between the hierarchy of nonzero responses and that of buffering structures and propose an algorithm to depict the hierarchy graph. We assume that CRNs do not have conserved quantities in the main text, but our results can be generalized to CRNs with conserved quantities in the Appendix.

## 2 The setting and review of structural sensitivity analysis

### 2.1 Setting

We label chemicals by *m* (*m* = 1, …, *M*) and reactions by *n* (*n* = 1, …, *N*), and consider a spatially homogeneous CRN [2, 24, 25, 26, 27, 5] consisting of the following reactions:

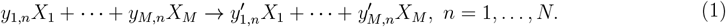

A state of the reaction system, specified by concentrations *x*_*m*_(*t*), obeys the differential equations

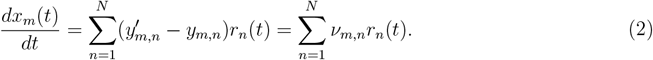

Here, the *M × N* matrix ***ν*** is called the stoichiometric matrix, defined as 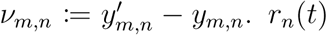 is the reaction rate (flux) of the reaction *n*. With ***x***(*t*) := (*x*_1_(*t*), …, *x*_*M*_ (*t*))^⊤^, ***r***(*t*) := (*r*_1_ (*t*), …, *r*_*N*_(*t*))^⊤^, (2) can be written as

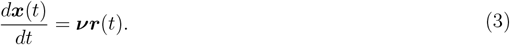

We assume that the flux function *r*_*n*_ depends on its reaction rate parameter *k*_*n*_, i.e.,

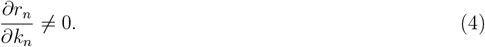

In metabolic networks, *k*_*n*_ can be the activitity or the amount of the enzyme catalyzing the reaction *n*. In addition, we assume that each flux function *r*_*n*_ is strictly increasing with respect to the concentrations of its substrates. We also account for regulations such as allosteric effects.

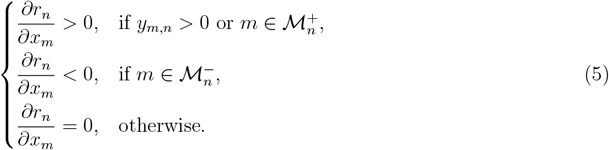

where 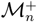 and 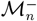 are the set of chemicals that are not a substrate of the reaction *n* but regulate the reaction positively and negatively, respectively. Popular kinetics, such as the mass-action 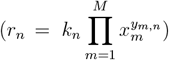 and the Michaelis-Menten kinetics (e.g. 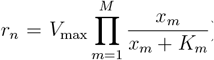), satisfy this condition. In the following, we do not assume specific forms of *r*_*n*_ except for the assumptions in (4) and (5).

### 2.2 Structural sensitivity analysis

We briefly review the structural sensitivity analysis [1, 2, 3]. This analysis allows us to determine qualitative changes in the steady-state concentration of each chemical *x*_*m*_ in response to changes in a reaction rate parameter *k*_*n*_ (sensitivities). We assume that the system has a steady state ***x***. At the steady state, the steady-state flux vector ***r*** satisfies ***νr*** = **0**, i.e., ***r*** ∈ ker ***ν***. We choose a basis for ker ***ν*** as 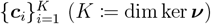. ***r*** can be written as a linear combination of 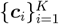;

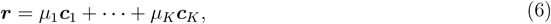

where *μ*_1_, …, *μ*_*K*_ ∈ ℝ.

Under the perturbation of *k*_*n*_→*k*_*n*_ + *δk*_*n*_, each flux changes from *r*_*j*_ to *r*_*j*_ + *δr*_*j*_. *δr*_*j*_ can be written as

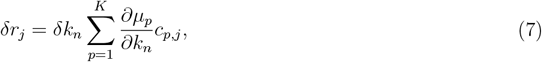

where *c*_*p,j*_ is the *j* th entry of ***c***_*p*_. At the same time, given that *r*_*j*_ depends on *k*_*j*_ and ***x***, i.e., *r*_*j*_ = *r*_*j*_ (*k*_*j*_, *x*_1_, …, *x*_*M*_), the total derivative of *δr*_*j*_ is given by

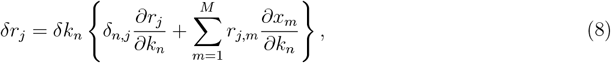

where *δ*_*n,j*_ is a Kronecker delta and 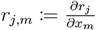.

From (7) and (8),

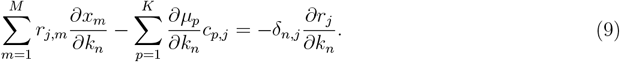

Let us define the matrices ***A*** ∈ ℝ^*N ×*(*M* +*K*)^ and 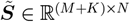at the steady state as follows.

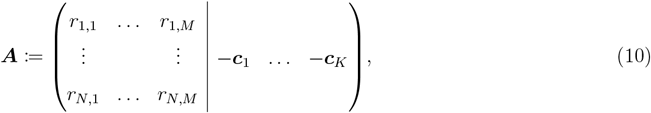

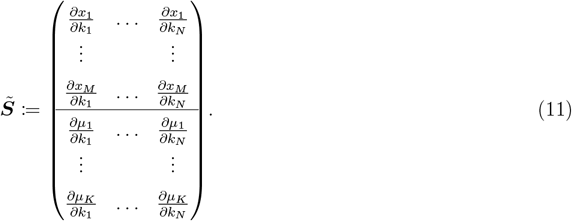

All entries of these matrices are partial derivatives evaluated at the steady state. The upper part of 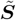 represents the rate of change in the steady-state concentration of each chemical with respect to each reaction rate parameter. The lower part of 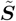 reflects the rate of change in the steady-state flux, which is given by (7).

From (9), we obtain

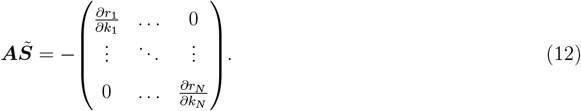

To present the key idea, we assume that ***A*** and 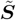 are square matrices, i.e., *M* + *K* = *N* . In this case, dim ker ***ν***^⊤^ = 0 holds, implying that there are no conserved quantities in the system (see Appendix for the general case). With (4) in mind, we obtain the following result.

#### Theorem 1.

[1] If ***A*** is invertible,

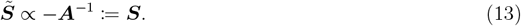

Here, ***X*** ∝ ***Y*** means the algebraic distribution of zero entries of matrices ***X*** and ***Y*** are the same. Using (5), the distribution of zero entries of ***A*** is algebraically determined from the network structure, so is that of 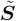. If *S*_*m,n*_ = 0 holds algebraically, the perturbation of *k*_*n*_ never affects the steady-state concentration of the chemical *m*, regardless of the values of *r*_*j,m*_ (≠ 0), in which case we say “the reaction *n* does not influence the chemical *m*”. If *S*_*m,n*_ ≠ 0 algebraically, there can exist some parameters or kinetics such that the perturbation of *k*_*n*_ affects the steady-state concentration of the chemical *m*, in which case we say “the reaction *n* influences the chemical *m*”. Overall, qualitative changes in the steady-state concentrations of chemicals in response to perturbation to each reaction rate parameter can be determined solely from the topology of a CRN. Note that if 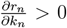 holds for *n* = 1, …, *N*, the signs of each entry in 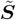 and those in ***S*** coincide, indicating that the signs of changes in the steady-state concentraions of each chemical can be determined from network topology.

#### Remark 1. *A* and *S* depend on the choice of the basis for ker *ν*

However, the sensitivity of the concentration of each chemical and flux at the steady state with respect to each reaction rate parameter *k*_*n*_ is independent of the choice of the basis, which we will prove in Lemma 1 (Section 3).

#### Definition 1. (Regularity of a CRN)

A CRN is called regular if it admits a steady state and the associated ***A*** is invertible. Throughout the paper, we assume the regularity so that ***A*** is invertible and the sensitivity ***S*** := −***A***^−1^ can be calculated [28].

### 2.3 Buffering structure

The concept of buffering structures derived from SSA is important for understanding, in terms of network topology, the extent to which the effects of changes in parameters propagate within a CRN [2, 3]. When a subnetwork of a regular CRN satisfies certain topological conditions, the subnetwork is called a buffering structure (as defined in Definition 2). It was mathematically demonstrated that the steady-state responses to the perturbation of a parameter within a buffering structure are confined within the buffering structure (as shown in Theorem 2, the law of localization). We briefly review the law of localization and buffering structure [2].

#### Definition 2. (Buffering structure)

[2] A subnetwork Γ = (𝔪_Γ_, 𝔯_Γ_) (𝔪_Γ_, 𝔯_Γ_ are subsets of chemicals and reactions, respectively) of a regular CRN is a buffering structure if it satisfies

1. None of the reaction rates of the reactions in 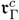 are dependent on the concentrations of chemicals within 𝔪_Γ_, i.e., 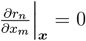 at all ***x***, 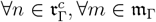 where 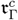 means the complement of 𝔯_Γ_ in all reactions in the CRN (Output-complete).
2. *λ*(Γ) = 0 with *λ*(Γ) = −|𝔪_Γ_| + |𝔯_Γ_| − *N* (𝔯_Γ_).

Here, |𝔪_Γ_| and |𝔯_Γ_| are the size of 𝔪_Γ_ and 𝔯_Γ_, respectively, *N* (𝔯_Γ_) is the number of stoichiometric cycles in 𝔯_Γ_. To be precise, *N* (𝔯_Γ_) := dim *{****x*** ∈ ker ***ν*** | supp ***x*** ⊂ 𝔯_Γ_*}*, where supp ***r*** := *{i* | *r*_*i*_ ≠ 0*}* for ***r*** ∈ ℝ^*N*^ . In the graphical representation of a CRN, if Γ is output-complete, no reaction arrows outside of Γ leave from the nodes (chemicals) inside Γ. It is important to note that the definition of a buffering structure does not depend on the choice of a basis for ker ***ν***.

Using Theorem 1 we can deduce Theorem 2, the proof of which was described in [2].

#### Theorem 2. (Law of localization)

[2] The steady-state chemical concentrations and reaction fluxes outside of a buffering structure Γ = (𝔪_Γ_, 𝔯_Γ_) do not change under any perturbation of the reaction rate parameters in 𝔯_Γ_.

Theorem 2 implies that when a subnetwork within a CRN satisfies certain topological conditions, the impact of a parameter perturbation within a buffering structure is restricted to that structure. A buffering structure introduces a novel concept in CRNs: a “regulatory module” can naturally arise from the topology of the network.

We illustrate SSA and buffering structures in an example network.

#### Example 1.

We consider a straight pathway, shown in Fig. 1A. The stoichiometric matrix is given by

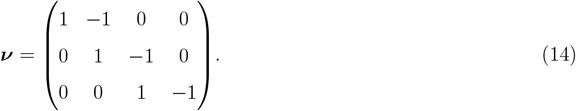

**Fig. 1:**
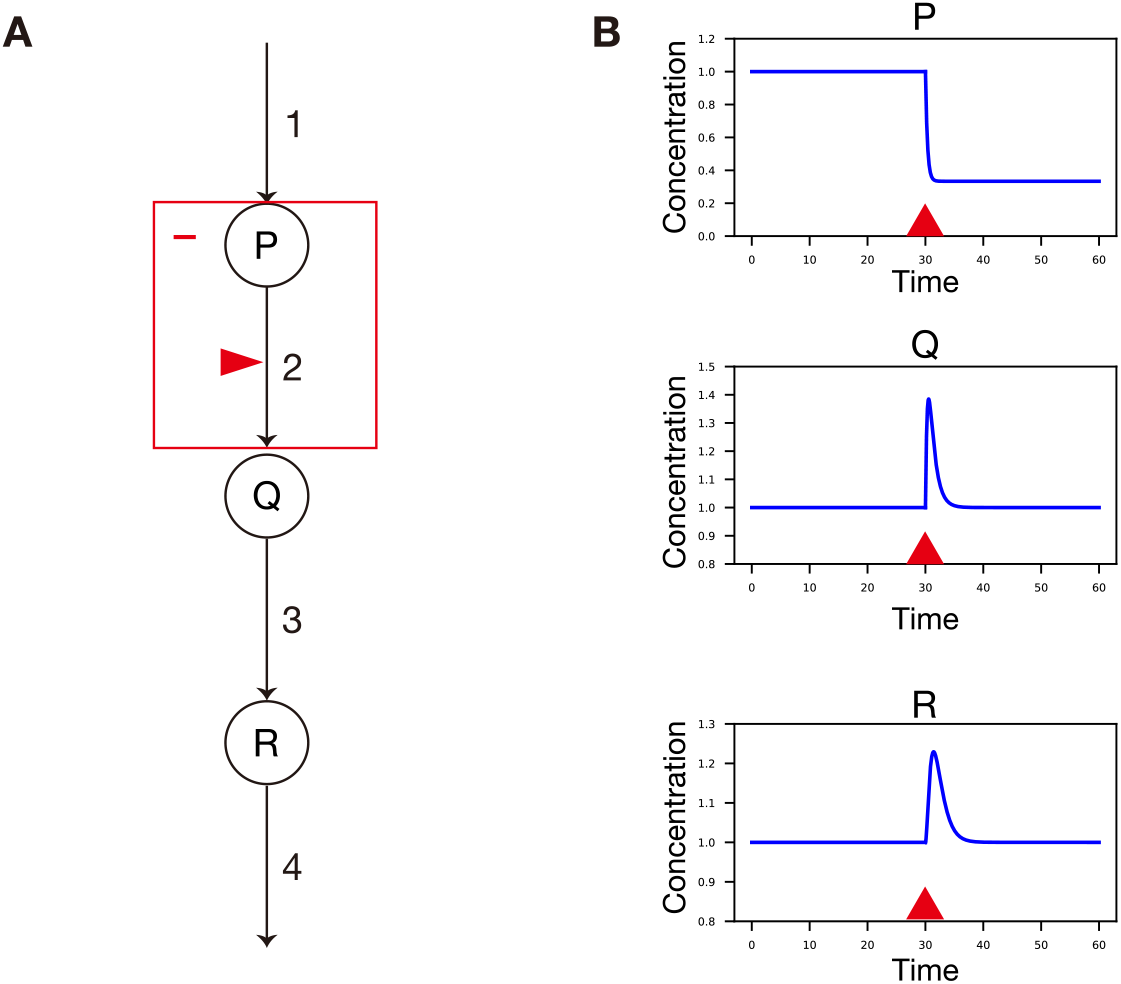
Analysis for a hypothetical network (Example 2). **A**, Graphical representation of a CRN comprising three chemicals (P,Q,R) and four reactions (1,2,3,4). Solid lines indicate chemical reactions. The subnetwork Γ = (*{P }, {*2*}*) is a buffering structure. **B**, The result of a numerical simulation in the network **A**. We assume ***r*** = (*k*_1_, *k*_2_*x*_*P*_, *k*_3_*x*_*Q*_, *k*_4_*x*_*R*_) with (*k*_1_, *k*_2_, *k*_3_, *k*_4_) = (1.0, 1.0, 1.0, 1.0). The dynamics are in the steady state at *t* = 30. At *t* = 30, *k*_2_ was increased by a factor of 2.

Since ***ν*** has a kernel vector ***c*** = (1, 1, 1, 1)^⊤^, the matrix **A** is

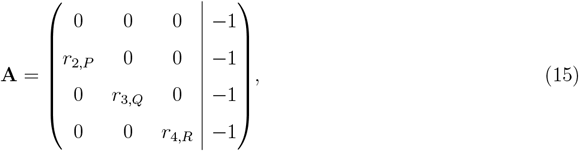

and the sensitivity is determined as

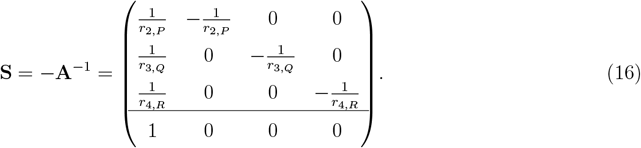

Since *r*_2,*P*_, *r*_3,*Q*_, *r*_4,*R*_ *>* 0 from (5), the distribution of nonzero entries in **S** can be determined. If 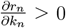, ∀*n*, the signs of changes in each chemical is given by

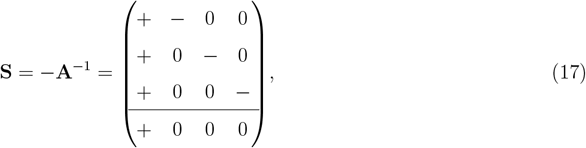

where + and − represent qualitative responses under perturbations associated with column indices. For example, from the second column, we can see that the increase in the reaction rate parameter of the reaction 2 results in a decrease in the steady-state concentration of the chemical P, but the steady-state concentrations of either Q or R are not affected (Fig. 1A,B). The subnetwork Γ = (*{P }, {*2*}*) is a buffering structure, because Γ is output-complete and *λ*(Γ) = −|𝔪_Γ_| + |𝔯_Γ_| − *N* (𝔯_Γ_) = −1 + 1 − 0 = 0 (Fig. 1A, red box). This explains why the steady-state concentrations of Q and R are insensitive to the perturbation to the reaction 2.

It was proved that the intersection or union of buffering structures is also a buffering structure [28].

#### Proposition 1.

Let Γ_1_ and Γ_2_ be buffering structures of a regular CRN (Γ_1_ ≠ Γ_2_). Then Γ_1_ ∪ Γ_2_ and Γ_1_ ∩ Γ_2_ are also buffering structures [28].

## 3 The main theorem

In this section, we present our main theorem: the inverse theorem of the law of localization. This theorem forms the basis for an efficient algorithm to identify buffering structures (Section 4). We first define a “response block”, which is an output-complete subnetwork with a confined response.

### Definition 3. (Response block)

A subnetwork Ψ = (𝔪_Ψ_, 𝔯_Ψ_) (𝔪_Γ_, 𝔯_Ψ_ are subsets of chemicals and reactions, respectively) of a regular CRN is a response block if it satisfies

1. Ψ is output-complete.
2. The steady-state concentrations of 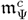 (chemicals outside of Ψ) do not change under the perturbation of either parameter in 𝔯_Ψ_.

### Remark 2.

The condition 2 of Definition 3 does not require that the steady-state reaction fluxes outside of Ψ remain unchanged under perturbations of reaction rate parameters in Ψ. However, this can be easily derived from the requirements of Definition 3. Indeed, if a subnetwork Ψ = (𝔪_Ψ_, 𝔯_Ψ_) is a response block, it is output-complete, which implies that reaction rates of 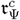 depend only on chemicals in 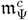. The steady-state concentrations of 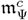 do not change under either perturbations of reaction rate parameters in Ψ, therefore, steady-state reaction rates of 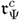 are not influenced by the perturbations.

From the law of localization (Theorem 2), a buffering structure is a response block. We will prove the main theorem of this paper: the inverse theorem of the law of localization (Theorem 3), which states that a response block is a buffering structure.

### Theorem 3. (Inverse theorem of law of localization)

Let Ψ be a response block of a regular CRN. Then, Ψ is a buffering structure.

From Theorem 2 and Theorem 3, we obtain the following proposition.

### Proposition 2. (The equivalence between a buffering structure and a response block)

Let Ψ be the subnetwork of a regular CRN. Then, the following are equivalent.

i. Ψ is a buffering structure.
ii. Ψ is a response block.

To prove Theorem 3, we will prove two lemmas.

### Lemma 1.

The sensitivities of the steady-state concentration of each chemical and flux with respect to reaction parameters *k*_*n*_ are indifferent to the choices of a basis for ker ***ν***.

### Proof of Lemma 1

Let *{****c***_*i*_*}*_*i*=1,…,*K*_ and 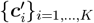 be two distinct bases for ker ***ν***. We define ***A*** and ***A***^*′*^ as

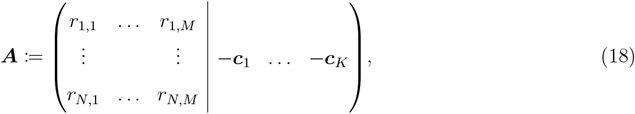

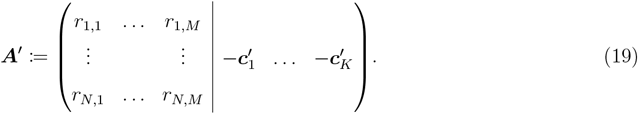

There exists an invertible change-of-basis matrix ***P*** ∈ ℝ^*K×K*^ such that 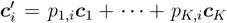 for *i* = 1, …, *K*. Using ***P***, we can rewrite ***A***^*′*^ as

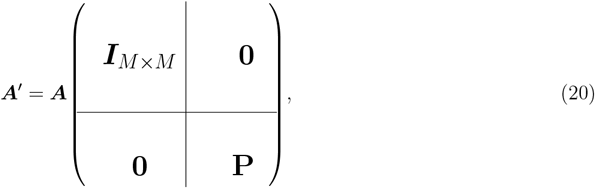

From this equation, the sensitivity matrix ***S***^*′*^ := −(***A***^*′*^)^−1^ can be written in the form

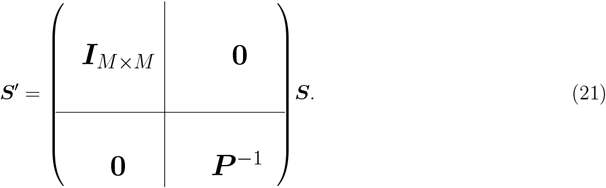

This equation shows that 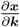, which is located in the first *M* rows of the sensitivity matrices, is the same for ***S*** and ***S***^*′*^. The sensitivity of each flux is given by (8) (since we are focusing on qualitative changes, we can assume 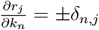), which is also the same between the two bases. □

Lemma 1 guarantees that the sensitivities of chemical concentrations fluxes with respect to reaction rate parameters are not affected by the choice of a basis for ker ***ν***, which allows us to choose any basis we prefer. The following lemma provides one way to choose a basis for ker ***ν***.

#### Lemma 2.

Let 𝔯_Γ_ be a subset of reactions in a CRN. We let 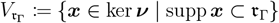. We also define 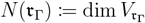 as previously defined. If dim ker ***ν*** ≥ 1, there exists a basis *{****f***_*i*_*}*_*i*=1,…,*K*_ for ker ***ν*** such that

i. 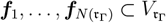
ii. 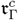 -projected 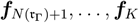 are linearly independent.

By 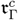 -projected ***f***, we mean 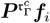, where 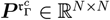 is a projection matrix satisfying

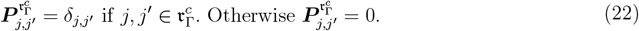

### Proof of Lemma 2

If *N* (𝔯_Γ_) = *K*, the statement is trivial. We will focus on the case of *N* (𝔯_Γ_) *< K*. If *N* (𝔯_Γ_) ≥ 1, we first choose a basis for 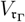 as 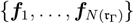, which satisfies condition (i). Then, we add some vectors to the basis vectors such that 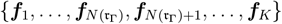 forms a basis for ker ***ν***. The set 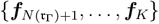 spans the complementary subspace of 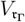, denoted by 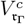 . (If *N* (𝔯_Γ_) = 0, we can choose an arbitrary basis for ker ***ν*** as *{****f***_*i*_*}*_*i*=1,…,*K*_. In this case, 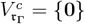.)

Let 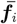 be the 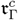 -projected ***f***_*i*_. We will prove that 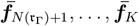 are linearly independent. Suppose on the contrary that 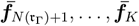 are linearly dependent. Then, there exists 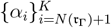 (at least one of 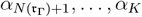 is nonzero) such that 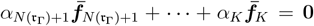. We let 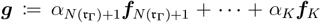. Because the 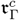 -projected ***g*** is **0**, we have ***g*** ∈ *V* (𝔯_Γ_). At the same time, ***g*** ∈ *V* (𝔯_Γ_)^*c*^ because ***g*** is written as a linear combination of 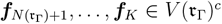. Therefore, ***g*** ∈ *V* (𝔯_Γ_) ∩ *V* (𝔯_Γ_)^*c*^ = *{***0***}*, i.e., ***g*** = **0**. This implies that 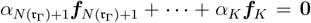, which contradicts the fact that 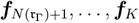 are linearly independent. This completes the proof of Lemma 2. □

### Proof of Theorem 3

Suppose a subnetwork Ψ = (𝔪_Ψ_, 𝔯_Ψ_) is a response block. Since Ψ is an output-complete subnetwork, it suffices to show that *λ*(Ψ) = 0.

We choose *{****f***_*i*_*}*_*i*=1,…,*K*_ for ker ***ν*** as shown in Lemma 2. As shown below, by collecting the indices associated with Ψ = (𝔪_Ψ_, 𝔯_Ψ_) into the upper-left corner, ***A*** can be represented as a block matrix with the lower-right corner being a zero matrix (Fig. 2, (Left)). The column indices in the upper left block consist of the chemicals in 𝔪_Ψ_ followed by 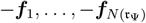, which are the basis vectors of 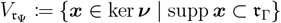. The row indices consist of the reactions in 𝔯_Ψ_. All entries in the green region in Fig. 2 (Left) are 0, because Ψ is output-complete. All entries in the blue region in Fig. 2 (Left) are 0, because supp ***f***_*i*_ ⊂ 𝔯_Ψ_ (*i* = 1, …, *N* (𝔯_Ψ_)). Since we are assuming Det **A** ≠ 0, the block associated with Ψ = (𝔪_Ψ_, 𝔯_Ψ_) in Fig. 2 (Left) is vertically long or square; i.e., *λ*(Ψ) = −|𝔪_Ψ_| + |𝔯_Ψ_| − *N* (𝔯_Ψ_) ≥ 0.

**Fig. 2:**
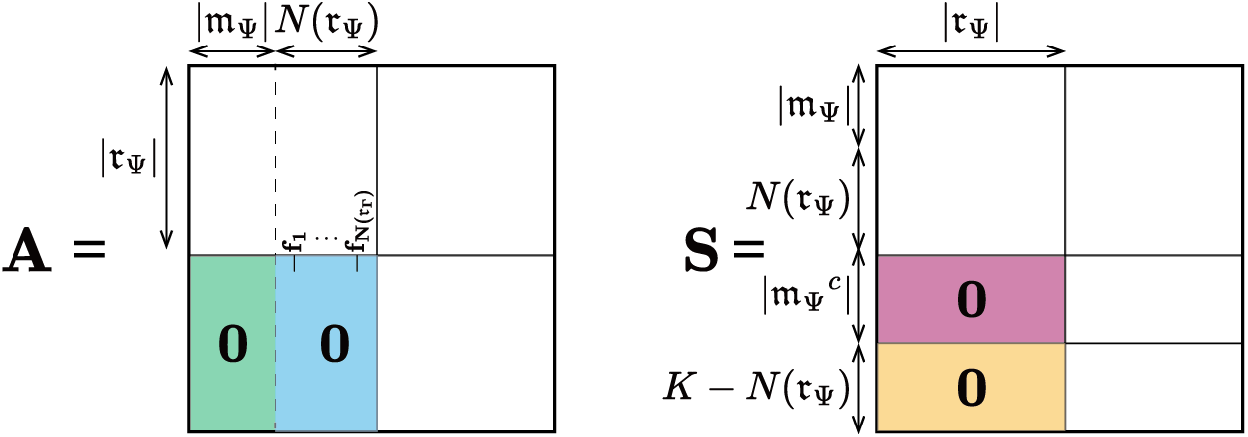
Schematic depiction of *A* and *S* when Ψ = (𝔪_Ψ_, 𝔯_Ψ_) is a response block. (Left) By collecting the indices associated with Ψ into the upper-left corner, ***A*** can be a block matrix in which the lower right is the zero matrix. (Right) The sensitivity matrix ***S*** is also proved to be a block matrix in which the lower right is the zero matrix. See proof of Theorem 3.

It remains to prove that *λ*(Ψ) ≤ 0. ***S*** = −***A***^−1^ is also in the block form in which the lower left part is the zero matrix, as we will prove below (Fig. 2 (Right)). First, all 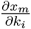 with 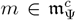 and *i* ∈ 𝔯_Ψ_, which appear in the red region in Fig. 2 (Right), vanish due to the assumption that the steady-state chemical concentrations in 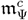 do not change under perturbations of reaction rate parameters in 𝔯_Ψ_. We next show that all 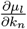, with 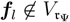 and *n* ∈ 𝔯_Ψ_, which appear in the orange region in Fig. 2 (Right), do not have nonzero entries. The change rate in the steady-state reaction flux vector in response to the perturbation to *k*_*n*_ is given by

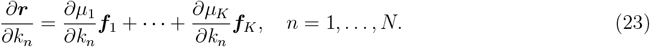

Recall that the reaction fluxes outside of a response block Ψ remain unchanged under either perturbations of reaction rate parameters in Ψ (Remark 2). This means that

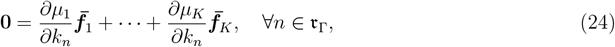

where 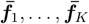 are 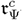 -projected ***f***_1_, …, ***f***_*K*_. From Lemma 2 (i), 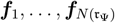 have nonzero values only in 𝔯_Ψ_, so 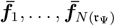 are all **0**. Thus we obtain

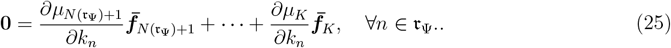

By Lemma 2, 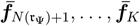 are linearly independent, which implies that

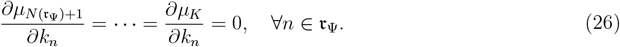

This completes the proof of the structure of ***S***. Because ***A*** is invertible, ***S*** is also invertible. This implies that the upper-left block in Fig. 2 (Right) is vertically long or square; i.e., |𝔪_Ψ_| + *N* (𝔯_Γ_) ≥ |𝔯_Ψ_|, thus *λ*(Ψ) ≥ 0, which completes the proof. □

## 4 The algorithm to find buffering structures

From Proposition 2, finding buffering structures is equivalent to finding response blocks. For each reaction *n*, we find the minimum buffering structure containing the reaction *n* through the following procedures (Algorithm 1).

We introduce some notations. For a reaction *n*, we let *M* (*n*) be the set of chemicals whose steady-state concentrations are affected by the perturbation to *k*_*n*_, i.e., 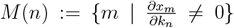. For a reaction set *𝒩*, we let *M* (*𝒩*) := ⋃ _*n*∈*𝒩*_ *M* (*n*). For a chemical *m*, we let *R*(*m*) be the set of reactions whose reaction rates are dependent on *m*, i.e., 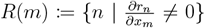. Intuitively, *R*(*m*) is the set of edges (reaction arrows) that leave the node (chemical) *m* in the graph. For a chemical set 𝔪, we let *R*(𝔪) := ⋃ _*m*∈𝔪_ *R*(*m*).

We first construct a subnetwork that contains only reaction *n*. Then, we add to the subnetwork *M* (*n*), whose steady-state concentrations are affected by the perturbation to *k*_*n*_. The subnetwork can be made output-complete by adding *R*(*M* (*n*)) *\ {n}*. From the transitivity property (Remark 3, below), the perturbations to the newly added reactions *R*(*M* (*n*)) *\ {n}* do not affect the steady-state concentration of any chemical outside the subnetwork. Hence, the resulting subnetwork *Y*_*n*_ is a response block and thus a buffering structure. It is clear that *Y*_*n*_ is the minimum buffering structure containing the reaction *n*. We iterate the procedure for *n* = 1, …, *N* and remove duplicates, obtaining 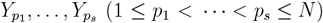.

### Remark 3.

If the system does not have the conserved quantities, the transitivity property for influences was proved [29]. If *r*_1_ influences *m*_1_, the reaction rate of *r*_2_ is dependent on *m*_1_, and the *r*_2_ influences *m*_2_, then *r*_2_ influences *m*_2_. In symbols,

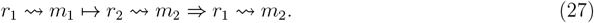

Here, *r* ⇝ *m* means that *m* influences *r* and *m* → *r* means that the reaction rate of *r* is dependent on the concentration of *m*.

### Algorithm 1 Finding all buffering structures (for CRNs without conserved quantities)

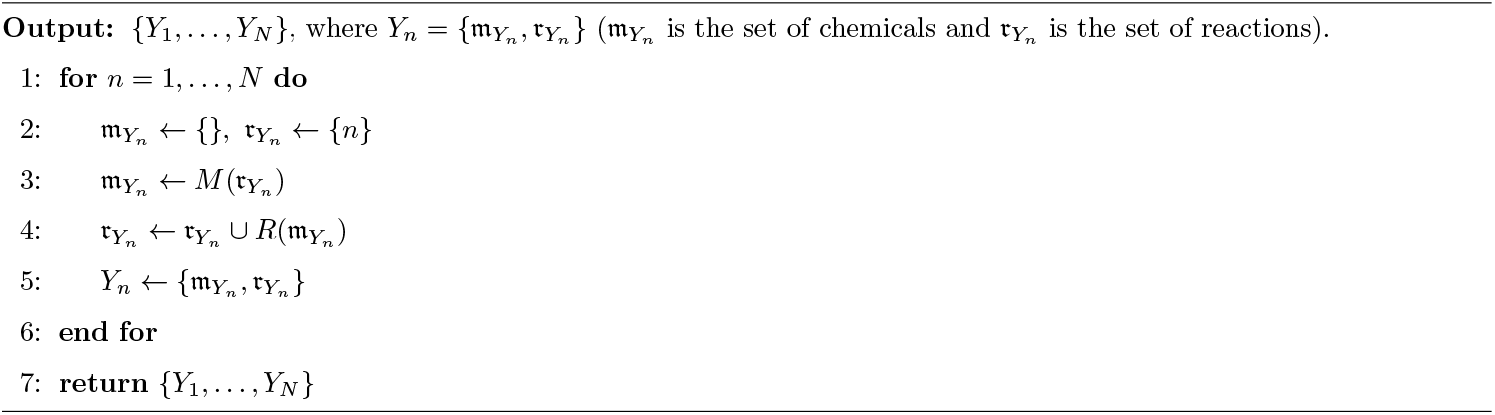

There exist buffering structures other than the aforementioned 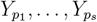. However, 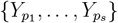 cover all buffering structures in a CRN in the sense that 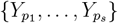 is the “minimum set of buffering structures” defined as follows.

### Definition 4. (Minimum set of buffering structures)

A set of buffering structures *{*Γ_1_, …, Γ_*s*_*}* is the “minimum set of buffering structures” of a regular CRN if

1. For any buffering structure Γ of the CRN, there exists 1 ≤ *n*_1_ *< … < n*_*m*_ ≤ *s* such that 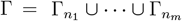.
2. Each Γ_*i*_ cannot be expressed as a union of smaller buffering structures, i.e., for any *i* (1 ≤ *i* ≤ *s*), there cannot exist nonempty buffering structures Γ*′* and Γ*′′* (Γ*′* ⊊ Γ_*i*_ and Γ*′′* ⊊ Γ_*i*_) such that Γ_*i*_ = Γ*′* ∪ Γ*′′*.

### Theorem 4.

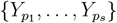 is the minimum set of buffering structures.

### Proof of Theorem 4

We first prove that 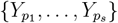 satisfies condition (1) of Definition 4. Suppose on the contrary that there exists a buffering structure Γ = (𝔪_Γ_, 𝔯_Γ_) that cannot be expressed as 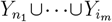. If Γ contains no reactions, |𝔯_Γ_| = |𝔪_Γ_| + *N* (𝔯_Γ_) = 0, hence |𝔪_Γ_| = 0 and Γ is empty. Therefore we only consider the case where Γ contains at least one reaction. Let *{n*_1_, …, *n*_*q*_*}* be the set of reactions in 𝔯_Γ_. For *i* = 1, …, *q*, Γ includes the mimimum buffering structure containing the reaction *n*_*i*_, hence we have Γ ⊃ *σ*, where 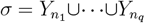. Because *σ* = (𝔪_*σ*_, 𝔯_*σ*_) is a buffering structure, |𝔪_*σ*_|−|𝔯_*σ*_|+*N* (𝔯_*σ*_) = 0. Since the reaction set in *σ* is the same as that in Γ, |𝔯_*σ*_| = |𝔯_Γ_| and *N* (𝔯_*σ*_) = *N* (𝔯_Γ_). Because |𝔪_Γ_|−|𝔯_Γ_| +*N* (𝔯_Γ_) = 0, we have |𝔪_*σ*_| = |𝔪_Γ_|. Since 𝔪_*σ*_ ⊂ 𝔪_Γ_, we have 𝔪_*σ*_ = 𝔪_Γ_ and thus *σ* = Γ, which means that 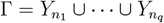, contradicting the assumption.

We finally prove that 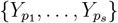 satisfies condition (2) of Definition 4. Suppose for some *i* ∈ ℕ (1 ≤ *i* ≤ *s*), there exists two non-empty buffering structures Γ*′* and Γ*′′* (Γ*′* ⊊ Γ and Γ*′′* ⊊ Γ) such that 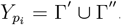. Since 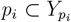, it holds that *p*_*i*_ ∈ Γ′ or *p*_*i*_ ∈ Γ″ . We can assume *p*_*i*_ ∈ Γ′ without loss of generality. Then, we have 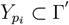. At the same time, since Γ*′* and Γ*′′* are nonempty, we have 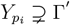, a contradiction. □

### Computational complexity

A naive algorithm to check whether a given subnetwork in a CRN conforms to the definition of BS (Definition 2) for all subnetworks has an exponential time complexity of *O*(2^*M*+*N*^). In contrast, the most time-consuming step of our proposed approach is the calculation of ***S*** = −***A***^−1^. Although symbolic calculation of the inverse matrix is infeasible for a large-sized matrix, the distribution of zero entries in the inverse of a sparse matrix can be estimated numerically [29]. We assign random values to *r*_*j,m*_ *(≠*0) appearing in ***A*** and numerically calculate ***A***^−1^. We repeat this process multiple times to determine the distribution of zero entries in ***S***. The time complexity of this procedure is *O*(*N* ^3^), which is much lower than the brute-force method.

We illustrate our algorithm with some example networks.

#### Example 2.

We consider a straight pathway, shown in Fig. 3A, which is the same as the pathway in Fig. 1A. As shown in (16), the sensitivity is given by

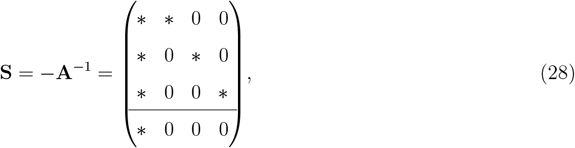

where * represents a nonzero response. We will exhaustively search for buffering structures in the CRN. Using the sensitivity matrix, we obtain *M* (*r*) as follows.

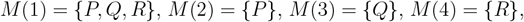

**Fig. 3:**
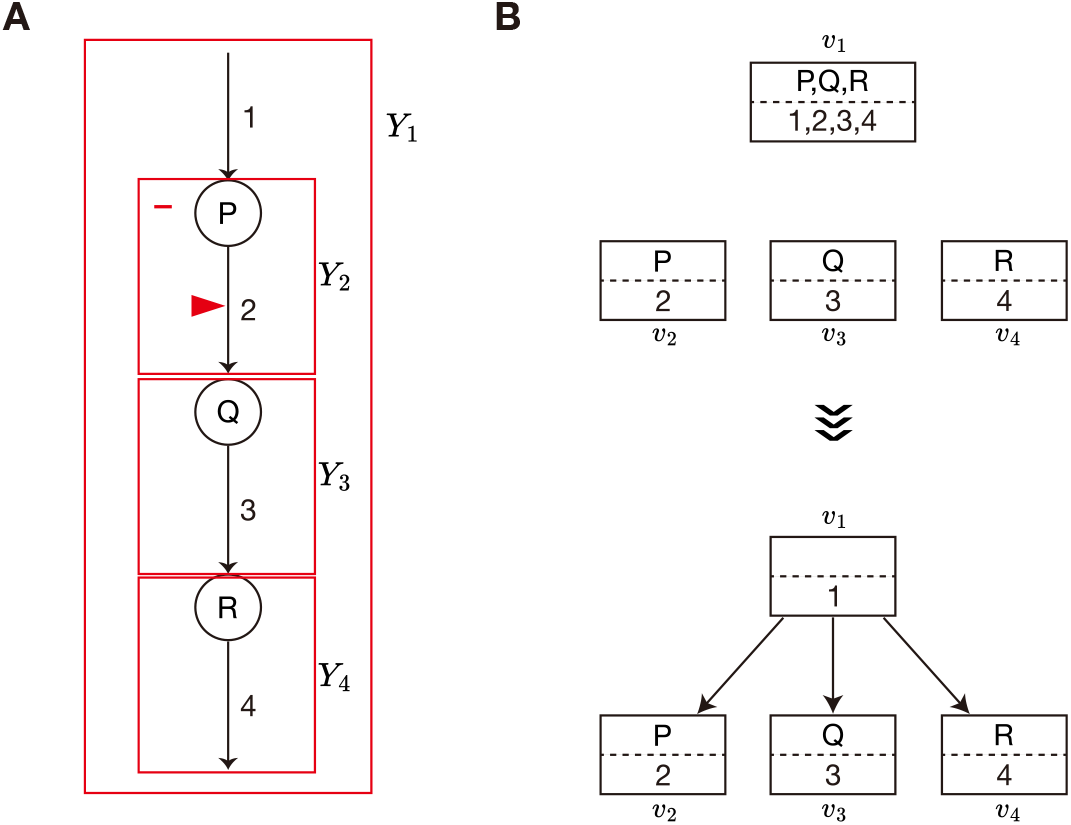
Analysis for a hypothetical network (Example 2). **A**, Graphical representation of a CRN with three chemicals (P,Q,R) and four reactions (1,2,3,4). Solid lines indicate chemical reactions. Each subnetwork enclosed by a red box (*Y*_*i*_) represents a minimum buffering structure containing the reaction *i*. **B**, Construction of the hierarchy graph. Initially, we construct a graph where each *Y*_*i*_ is assigned to a node *v*_*i*_. Subsequently, we build the hierarchy graph such that modulating the enzyme activity of reactions within a square box can lead to nonzero responses in the chemicals within that box and those in the lower boxes, while leaving the other chemicals unaffected.

We also obtain *R*(*m*), which is the set of edges (reaction arrows) that leave the node (chemical) *m* in the graph.

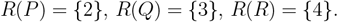

Since this system does not have conserved quantities, the minimum set of buffering structures is given by Algorithm 1 (Fig. 3A).

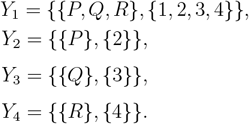

#### Example 3.

We consider a hypothetical pathway, shown in Fig. 4A. The matrix **A** is

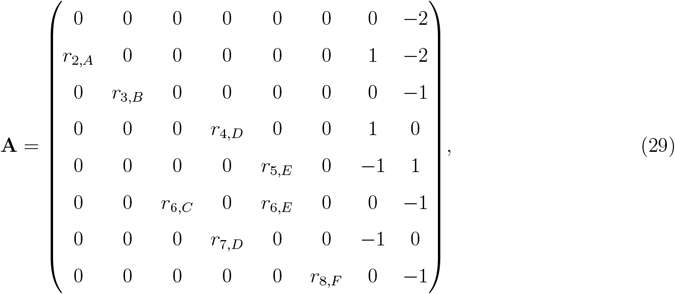

and the sensitivity is determined as

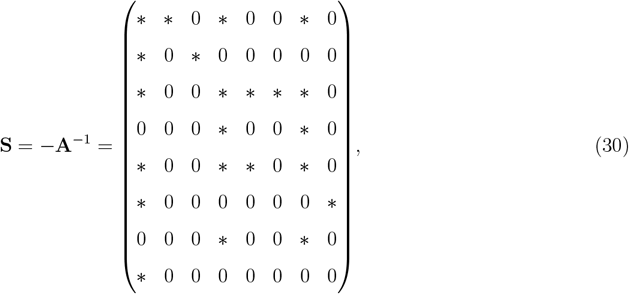

where * represents a nonzero response. Using the sensitivity matrix, we obtain *M* (*r*) as follows.

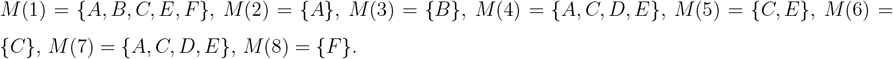

**Fig. 4:**
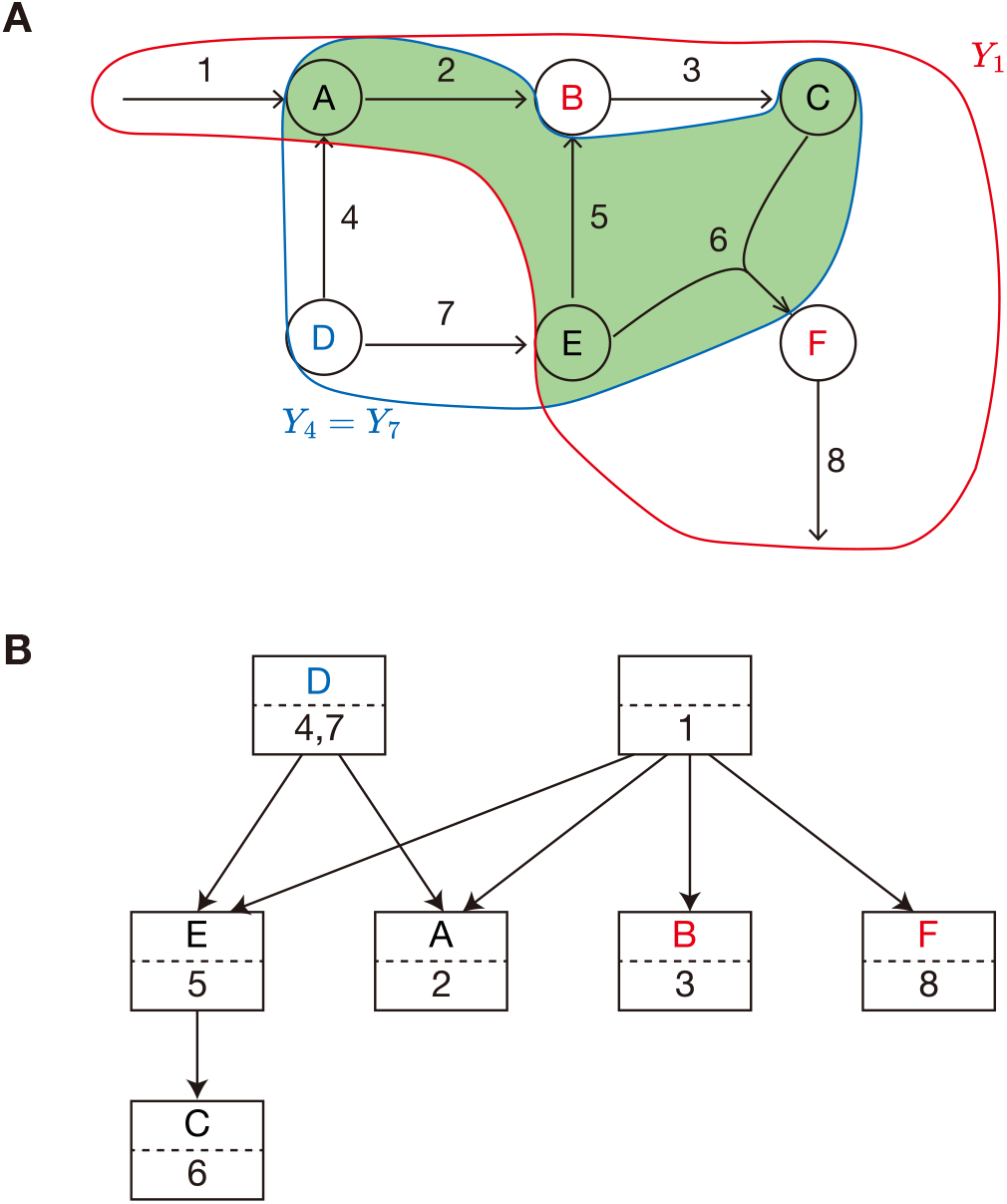
Analysis for a hypothetical network (Example 3). **A**, Graphical representation of a CRN with 6 chemicals (A,B,C,D,E,F) and 8 reactions (1, …, 8). Solid lines indicate chemical reactions. *Y*_1_ is shown in red, and *Y*_4_ = *Y*_7_ is shown in blue. The intersection of these two is colored in green, which is also a buffering structure. **B**, The hierarchy graph. Modulating the enzyme activity of reactions within a square box leads to nonzero responses in the chemicals within that box and those in the lower boxes, leaving the other chemicals unaffected.

We also obtain *R*(*m*), which is the set of edges (reaction arrows) that leave the node (chemical) *m* in the graph.

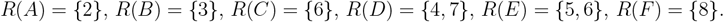

Since this system does not have conserved quantities, the minimum set of buffering structures is given by Algorithm 1 (Fig. 4A).

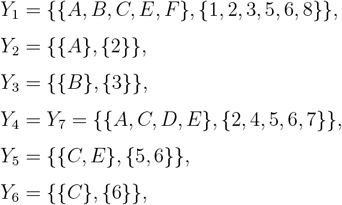

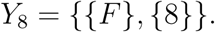

Parameters that influence D and those that affect B (or F) are disjoint, hence D and B (or F) are independently regulated. This is explained by the presence of two buffering structures. B and F are part of *Y*_1_ (Fig. 4A red), hence the reactions in *Y*_1_ influences only chemicals within *Y*_1_, including B and F. D is part of *Y*_4_ = *Y*_7_ (Fig. 4A blue), hence the reactions in *Y*_4_ = *Y*_7_ influences only chemicals within *Y*_4_ = *Y*_7_, including D. These two buffering structures have an intersection (Fig. 4A green), including the reactions 2, 5, and 6. From Proposition 1, the intersection is also a buffering structure, indicating that the reactions in the intersection do not influence B, D, or F. Thus, the reactions influencing D and those affecting B (or F) are disjoint. This is an example where multiple chemicals in a single connected CRN are independently regulated through buffering structures.

## 5 Hierarchy graph

We previously found that the nonzero response patterns under perturbations of different parameters can exhibit inclusion relations among them, i.e., exhibit hierarchical structures [2]. Based on the arguments in Section 4, we will show that the hierarchy among nonzero responses is equivalent to the hierarchy of buffering structures.

We are able to construct a hierarchy graph of nonzero response patterns in the following way. We obtain 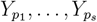 in the same way as Section 4. Each 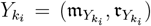 represents a perturbation-response relationship, i.e., perturbations to parameters in 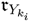 are confined to 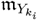 . In the following way, we construct the graph 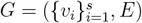 representing the inclusion relationship of 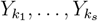.

1. We initially prepare a set of nodes 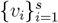. 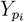 is assigned to *v*_*i*_. The edge set *E* is empty at this time.
2. For any pair of *i* and *j* (1 ≤ *i, j* ≤ *s*), we add an edge 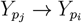 if 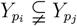.
3. For any pair of *i* and *j* (1 ≤ *i, j* ≤ *s*), we remove an edge 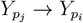 if there exists the path from 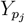 to 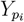 of length greater than or equal to 2. The graph *G* is a directed acyclic graph (If 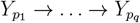 forms a cycle, 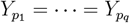, which contradicts the fact that duplicates in 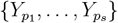 are removed).
4. We let 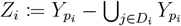, where *D*_*i*_ is the union of nodes that are downstream of *v*_*i*_. We change the assignment of *v*_*i*_ from 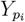 to *Z*_*i*_.

The resulting hierarchy graph *G* represents the inclusion relationship of nonzero response patterns. When a reaction rate in a node is perturbed, nonzero responses are confined to the chemicals in the node plus those in the lower nodes. From Theorem 3, each node *v*_*i*_ plus its downstream nodes *D*_*i*_ corresponds to a buffering structure.

### Theorem 5.

In the hierarchy graph *G*, every chemical and reaction appears exactly once.

### Proof of Theorem 5

Because *Y*_*i*_ contains reaction *i*, every reaction appears at least once. From the regularity of the matrix ***A, S*** is invertible, thus, every chemical is influenced by at least one reaction. Hence, all chemicals appear in *G* more than once.

Suppose *G* has two distinct nodes *v*_*i*_ and *v*_*j*_ such that *Z*_*i*_ ∩ *Z*_*j*_ ≠ ∅. We pick *e* ∈ *Z*_*i*_ ∩ *Z*_*j*_ (where *e* can be either a chemical or a reaction). Since 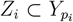 and 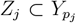, we have 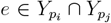. Because both 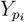 and 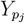 are buffering structures, their intersection 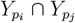 is also a buffering structure by Proposition 1. Thus, according to Theorem 4, there exist buffering structures 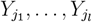 such that 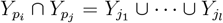. There exists *k* (1 ≤ *k* ≤ *l*) such that 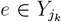 . Since 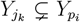, the node *v*_*k*_ is located downstream of *v*_*i*_, implying that 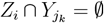. However, this contradicts the assumption that *e* ∈ *Z*_*i*_ and 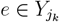. This completes the proof of Theorem 5. □

We illustrated our results in the example networks (Fig. 3B and Fig. 4B). Hierarchy graphs clearly visualize not only the nonzero response patterns but also the inclusion relationships between buffering structures.

## 6 Discussion

We previously demonstrated that the response of a CRN to the modulations of parameters can be determined solely from the local structure of the network [2, 3]. When a subnetwork within a CRN satisfies specific topological conditions, it is denoted as a “buffering structure”. The influence of a parameter perturbation within a buffering structure is confined within it (the “law of localization”). A buffering structure represents a novel concept in CRNs: a “regulatory module” can naturally arise from the topology of the network.

In previous studies [2, 3], an ad-hoc method was employed where buffering structures are identified by searching for an output-complete subnetworks that shows confined responses, based on SSA. However, it remained unclear whether all subnetworks exhibiting these properties satisfy the topological conditions of buffering structures. Our present study proves that an output-complete subnetwork displaying a finite response range is always a buffering structure. This shows that buffering structures can be computed without redundancy by searching for output-complete subnetworks that exhibit confined responses. A naive algorithm to check whether, for all subnetworks, a given subnetwork in a CRN conforms to the definition of a buffering structure, i.e., exhibits output-completeness and has an index of zero, has an exponential time complexity. In contrast, our proposed approach has polynomial time complexity. The existence of a hierarchy among the nonzero responses to parameter perturbations has been hinted at in prior work. However, the connection between this hierarchy of nonzero responses and that of buffering structures remained elusive. Our research elucidates that the hierarchy among nonzero responses is equivalent to the hierarchy of buffering structures. Furthermore, we propose a concrete algorithm to depict this hierarchy graphically through the computation of buffering structures.

This method can be applied to any CRN as long as the ***A*** matrix is invertible. Our approach equips researchers to explore buffering structures across various biological CRNs, including metabolic systems and signal transduction networks. This will clarify how different physiological functions emerging from a single CRN can be separately regulated.

In addition to sensitivities, the qualitative change (i.e., plasticity) of behaviors is another important aspect of biological systems. Mathematically, these plastic behaviors can be studied within the framework of bifurcation theory of dynamical systems. Previously, two of the authors of this study introduced a method to study bifurcations of CRNs based solely on network topology, termed ‘Structural Bifurcation Analysis’ (SBA) [30, 31]. SBA was developed upon an equivalence between the Jacobian matrix ***J*** of a reaction system and the augmented matrix ***A***, enabling bifurcation analysis based on network information. An important step in SBA involves decomposing a CRN according to the inclusion relationship among buffering structures. Our algorithm presented in this study for enumerating buffering structures and constructing their hierarchy will be useful for executing SBA.

The inverse theorem offers the potential utility for correcting the network information when combined with perturbation experiments. A vast amount of information about metabolic networks is available in databases [32, 33, 34, 35, 36, 37], but the information of networks in these databases might be still incomplete: there might exist unidentified reactions or regulations. By perturbing metabolic enzymes and experimentally measuring the responses, it will be possible to identify the subnetwork that exhibits a confined response range. The index of such a subnetwork should be zero according to our inverse theorem. If the index calculated from the database network is not zero, modifications should be made to the network to ensure that the index is zero. By employing this strategy, it may be possible in the future to refine the metabolic networks in databases.

From a medical point of view, our algorithm for identifying buffering structures might be useful for elucidating the mechanisms underlying drug resistance. Drugs designed to treat diseases, such as cancer, operate by inhibiting enzyme activity within CRNs. The extent of the molecular response induced by drugs could be defined by the buffering structure. It is plausible that, as cancer progresses, the structure of the CRN changes, leading to changes in buffering structures. If buffering structures narrow down, the effects of drug could get weaker, thereby inducing drug resistance. By comparing the buffering structures of normal tissues and those of cancerous CRNs, we may be able to identify the CRN changes that are key to drug resistance.

Hirono et al. proved the inverse theorem independently of us [22]. In our study, we not only proved the inverse theorem but also clarified the relationship with the hierarchy of nonzero responses and that of buffering structures.

One limitation of this study is that we have only discussed the case where the system is regular, i.e., the ***A*** matrix is invertible. When the system is not regular, we conjecture that structurally stable fixed points may not exist. If so, the ***A*** matrix in the actual biological CRNs should be invertible given that structurally stable fixed points are likely to exist. However, we observed that the ***A*** matrix calculated from the information in CRN databases is sometimes singular. One possibility is that network information within these databases may contain missing reactions or regulations. It is thus promising to develop a method to identify subnetworks that are predicted to be robust buffering structures even under possible network modifications.

In summary, we have developed an efficient method for exhaustively identifying buffering structures in any given CRN. This method is expected to lead to a better understanding of the basis for the independent regulation of different functions arising from a single connected CRN.

## 7 Code availability

The Python implementation of ibuffpy is available at https://github.com/hishidagit/SSApy.

## 8 Acknowledgements

We thank Yong Jin and Masato Ishikawa for their helpful discussions and comments. This research was supported by the CREST program (grant no. JPMJCR1922) of the Japan Science and Technology Agency (JST) (http://www.jst.go.jp/EN/index.html), Grant-in-Aid for Scientific Research on Innovative Areas (grant no. 19H05670), Joint Usage/Research Center program of Institute for Life and Medical Sciences Kyoto University. YY was supported by JSPS KAKENHI (Grant No. 23K14156). TO was supported by JSPS KAKENHI (Grant No. 22K03453, 22K06347) and the RIKEN iTHEMS Program. AH was supported by JSPS KAKENHI (Grant No. 23KJ1324).

## 9 Author contributions

Y.Y. designed the study, provided mathematical proofs for the theorems, and wrote the manuscript. A.H. implemented the algorithm, and Y.Y also contribute to the implementation. T.O and A.M. contributed to the supervision of the study and the writing of the manuscript. Y.Y. wrote the manuscript with assistance from the other authors.

## Appendix A CRNs with conserved quantities

### Appendix A.1 Structural sensitivity analysis with conserved quantities

In the main text, we assumed that the system does not have the conserved quantities, i.e., dim ker ***ν***^⊤^ = 0, and thus *M* + *K* = *N* . In [3], SSA is generalized to CRNs with conserved quantities, which allows us to study any CRNs. We denote a basis of ker ***ν***^⊤^ (the cokernel basis) as 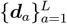, where *L* := dim ker ***ν***^⊤^. The quantity 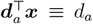 remains constant throughout the dynamics (a conserved quantity), since 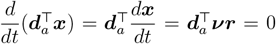. In the presence of conserved quantities, steady-state concentrations and fluxes are affected not only by reaction rate parameters but also by the initial values of conserved quantities. Therefore, in this case, there are two types of perturbations; the perturbation of the reaction rate parameter *k*_*n*_ and that of the conserved quantity *d*_*a*_. To treat two types of perturbations in a unified way, we introduce generalized parameters *J*_*i*_ (*i* = 1, …, *N* + *L*) as

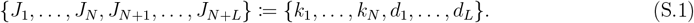

We generalize the definition of ***A*** and 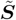 as follows.

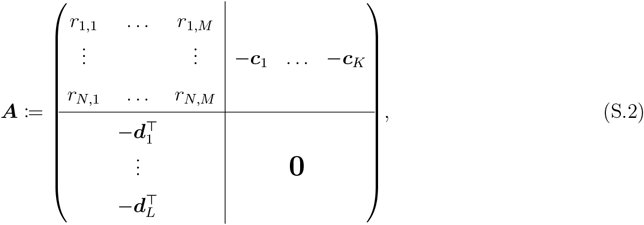

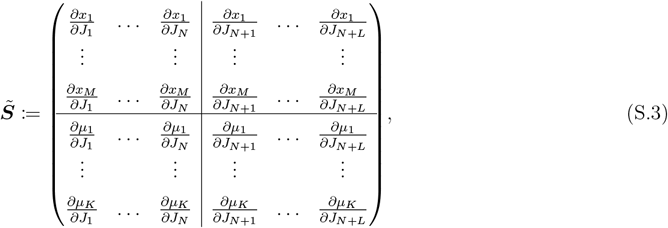

(12) is generalized to

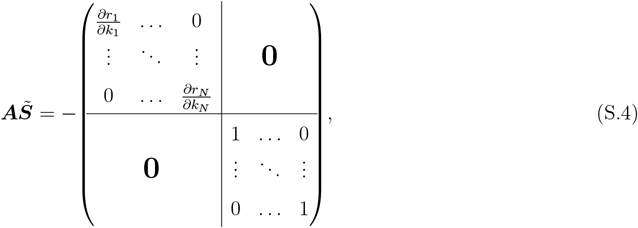

where 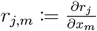. All partial derivatives are evaluated at the steady state.

With (4) in mind, Theorem 1 is generalized to Theorem S1.

#### Theorem S1.

If ***A*** is invertible,

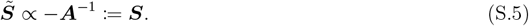

#### Remark S1. (The selection of the cokernel basis)

The values of 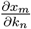 are independent of the choice of bases for ker ***ν*** and ker ***ν***^*T*^, which is proved in much the same way as Lemma 1. Hence, when our focus is solely on the effects of perturbations to the reaction rate parameters, we have the flexibility to choose the bases as we prefer. However, if our interest lies in the effects of perturbations to the conserved quantities, it is crucial to carefully select the basis for ker ***ν***^⊤^. If supp ***d***_*a*_ ⊂ supp ***d***_*b*_ for some *a, b* (1 ≤ *a* ≠ *b* ≤ *L*), the perturbation of *d*_*a*_ can affect *d*_*b*_, in which case the partial derivative 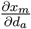 is difficult to interpret. To avoid choosing such a basis, it is recommended to find a basis by transforming ***ν***^⊤^ into its reduced row-echelon form (RREF). When finding buffering structures, choosing the cokernel basis requires additional caution (See also Appendix A.4.2).

### Appendix A.2 Buffering structure

We briefly review the law of localization and buffering structure [3].

#### Definition S1. (Generalized definition of a buffering structure)

[3] A subnetwork Γ = (𝔪_Γ_, 𝔯_Γ_) (𝔪_Γ_, 𝔯_Γ_ are subsets of chemicals and reactions, respectively) of a regular CRN is a buffering structure if it satisfies

1. None of the reaction rates of the reactions in 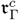 are dependent on the concentrations of chemicals within 𝔪_Γ_, i.e., 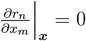 at all ***x***, 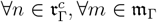, where 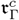 means the complement of 𝔯_Γ_ in all reactions in the CRN (Output-complete).
2. *λ*(Γ) = 0 with *λ*(Γ) = −|𝔪_Γ_| + |𝔯_Γ_| − *N* (𝔯_Γ_) + *N*_*c*_(𝔪_Γ_).

Here, |𝔪_Γ_| and |𝔯_Γ_| are the size of 𝔪_Γ_ and 𝔯_Γ_, respectively, *N* (𝔯_Γ_) is the number of stoichiometric cycles in 𝔯_Γ_. To be precise, *N* (𝔯_Γ_) := dim *{****x*** ∈ ker ***ν*** | supp ***x*** ⊂ 𝔯_Γ_*}*, where supp ***r*** := *{i* | *x*_*i*_ ≠ 0*}* for ***r*** ∈ ℝ^*N*^ . *N*_*c*_(𝔪_Γ_) is the number of independent conserved quantities containing at least one element in 𝔪_Γ_. To be precise, 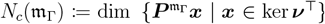, where 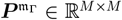 is a projection matrix onto the space associated with 𝔪_Γ_ defined as

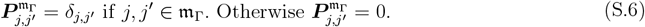

Note that the definition of a buffering structure is indifferent to the choice of bases for ker ***ν*** or ker ***ν***^⊤^.

From Theorem S1 we can deduce Theorem S2.

#### Theorem S2. (Law of localization)

[3] We consider a regular CRN with conserved quantities. We choose a basis for ker ***ν***^⊤^ as 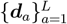 in the following way. First, we start with the basis vectors for 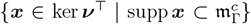. Then, we repeatedly add a vector not in the span of the vectors to the set until it spans ker ***ν***^⊤^. If a subnetwork Γ = (𝔪_Γ_, 𝔯_Γ_) (𝔪_Γ_, 𝔯_Γ_ are subsets of chemicals and reactions, respectively) is a buffering structure, then the following two hold.

1. For any reaction *n* ∈ 𝔯_Γ_ and for any chemical 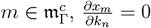 holds.
2. For any *a* satisfying supp ***d***_*a*_ ∩ 𝔪_Γ_ ≠ ∅ and for any chemical 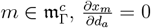 holds.

Theorem S2 states that the steady-state chemical concentrations outside of a buffering structure do not change under perturbations to reaction rate parameters or conserved quantities within the buffering structure. Since Γ is output-complete, the steady-state reaction fluxes outside of a buffering structure do not change under these perturbations either, as discussed in Remark 2.

### Appendix A.3 The inverse theorem

We provide a generalized definition of a response block.

#### Definition S2. (General definition of a response block)

In the case of a regular CRN with dim ker ***ν***^⊤^ *>* 0, we choose a basis of ker ***ν***^⊤^ as 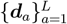. We define a subnetwork Ψ = (𝔪_Ψ_, 𝔯_Ψ_) (𝔪_Ψ_, 𝔯_Ψ_ represent subsets of chemicals and reactions, respectively) as a response block under the cokernel basis 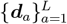 if it meets the following criteria:

1. Ψ is output-complete.
2a. For any reaction *n* ∈ 𝔯_Ψ_ and any chemical 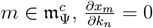.
2b. For any *a* such that supp ***d***_*a*_ ∩ 𝔪_Ψ_ ≠ ∅ and any chemical 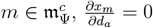.

Note that the definition of a response block depends on the choice of a basis of ker ***ν***^⊤^, whereas a buffering structure is defined independently of the choice. According to the law of localization (Theorem S2), a buffering structure is a response block for some choice of the basis for ker ***ν***^⊤^. It can be proved that the inverse theorem of the law of localization holds, which states that if a subnetwork is a response block under some choice of the basis for ker ***ν***^⊤^, the subnetwork is a buffering structure.

#### Theorem S3. (Generalized inverse theorem of the law of localization)

We consider a regular CRN. If dim ker ***ν***^⊤^ *>* 0, we choose a basis for ker ***ν***^⊤^ as 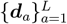. Let Ψ be a response block under the cokernel basis 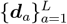. Then, Ψ is a buffering structure.

#### Proof of Theorem S3

The proof is almost the same as that of Theorem 3. Briefly, by appropriately choosing bases for ker ***ν*** and ker ***ν***^*T*^ and the orderings of the indices of ***A, A*** becomes a block matrix in which the lower right is the zero matrix (Fig. 5 (Left)). Since ***A*** is invertible, the left upper part of the block matrix is horizontally long or square, i.e., *λ*(Γ) = −|𝔪_Γ_| + |𝔯_Γ_| − *N* (𝔯_Γ_) + *N*_*c*_(𝔪_Γ_) ≥ 0. In addition, ***S*** becomes a block matrix in which the lower right is the zero matrix, which can be proved in much the same way as the proof of Theorem 3 (Fig. 5 (Right)). Since ***S*** is invertible, the left upper part of the block matrix is horizontally long or square, i.e., *λ*(Γ) = −|𝔪_Γ_| + |𝔯_Γ_| − *N* (𝔯_Γ_) + *N*_*c*_(𝔪_Γ_) ≤ 0. Thus, we obtain *λ*(Γ) = 0. Since Ψ is output-complete, Ψ is a buffering structure. □

**Fig. 5:**
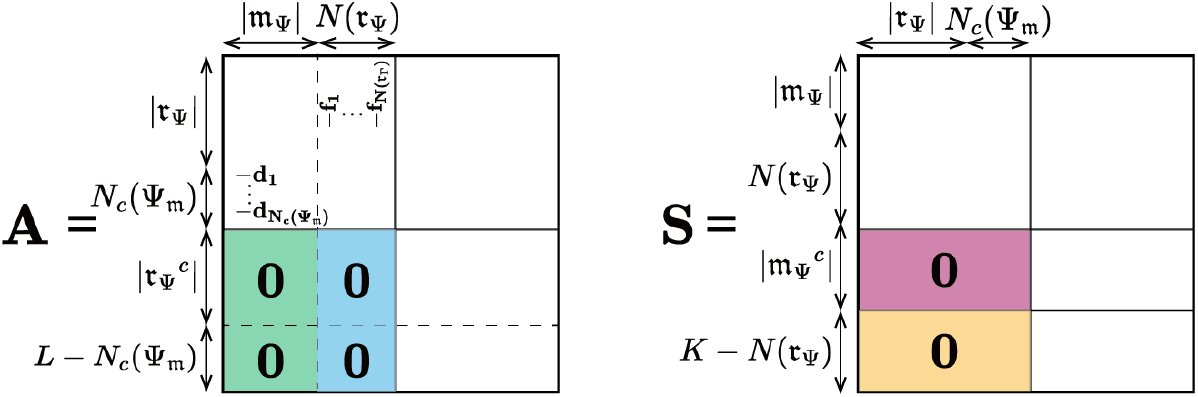
Schematic depiction of *A* and *S* when Ψ = (𝔪_Ψ_, 𝔯_Ψ_) is a response block. (Left) By collecting the indices associated with Ψ into the upper-left corner, ***A*** can be a block matrix in which the lower right is the zero matrix. (Right) ***S*** is also proved to a block matrix in which the lower right is the zero matrix. See proof of Theorem S3.

From Theorem S2 and Theorem S3, we obtain the following proposition.

##### Proposition S1. (The equivalence between a buffering structure and a response block)

Let Ψ be the subnetwork of a regular CRN. The following are equivalent.

1. Ψ is a buffering structure.
2. Ψ is a response block for some choice of the basis for ker ***ν***^⊤^.

### Appendix A.4 The algorithm to find buffering structures

#### Appendix A.4.1 Algorithm

We generalize the algorithm to find buffering structures in CRNs with conserved quantities. Here, we describe the algorithm to find response blocks under the specific choice of the basis for ker ***ν***^⊤^, denoted by 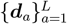 . How we should choose the basis is discussed in Appendix A.4.2. We define *{J*_1_, …, *J*_*N*_, *J*_*N*+1_, …, *J*_*N*+*L*_*}* := *{k*_1_, …, *k*_*N*_, *d*_1_, …, *d*_*L*_*}* as shown in (S.1). For each reaction rate parameter or conserved quantity *n* (1 ≤ *n* ≤ *N* + *L*), we find the minimum buffering structure containing parameter *n* through the following procedures (Algorithm 2).

Similar to the notations in the main text, we introduce some notations. For each parameter *n* (*n* = 1, …, *N* + *L*), which corresponds to either the reaction rate parameter or conserved quantity, we let *M* (*n*) be the set of chemicals whose steady-state concentrations are affected by the perturbation to *n*. For a reaction set *𝒩*, we let *M* (*𝒩*) := ⋃ _*n*∈*𝒩*_ *M* (*n*). For a chemical *m*, we let *R*(*m*) be the set of reactions whose reaction rates are dependent on *m*, i.e., 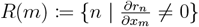. We also let *R*_*q*_(*m*) be the set of conserved quantities that contain *m*, i.e., *R*_*q*_(*m*) := *{d*_*N*+*a*_ | *m* ∈ supp ***d***_*a*_*}*. For a chemical set 𝔪, we let *R*(𝔪) := ⋃ _*m*∈𝔪_ *R*(*m*) and *R*_*q*_(𝔪) := ⋃_*m*∈𝔪_ *R*_*q*_(*m*).

We begin by constructing a subnetwork that contains only parameter *n*. Then, we add *M* (*n*) to the subnetwork. To make the subnetwork output-complete, we add *R*(*M* (*n*)) to the subnetwork. We also add *R*_*q*_(*M* (*n*)) to the subnetwork. Again, we add chemicals that are influenced by the newly added parameters. The subnetwork can be made output-complete by adding some reactions. We also add conserved quantities that contain at least one of the newly added chemicals. We repeat this procedure until there are no reactions or conserved quantities to be added. The resulting subnetwork *Y*_*n*_ is output-complete and the effects of the perturbations to the reaction rate parameters or conserved quantities in *Y*_*n*_ are confined inside *Y*_*n*_, indicating that *Y*_*n*_ is a response block under the basis 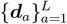. . According to Theorem S3, *Y*_*n*_ is a buffering structure. We iterate the procedure for *n* = 1, …, *N* + *L* and remove duplicates, obtaining 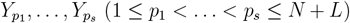. This algorithm stops in a finite number of steps since the number of chemicals plus the number reactions in a network is finite.

##### Algorithm 2 Finding all buffering structures

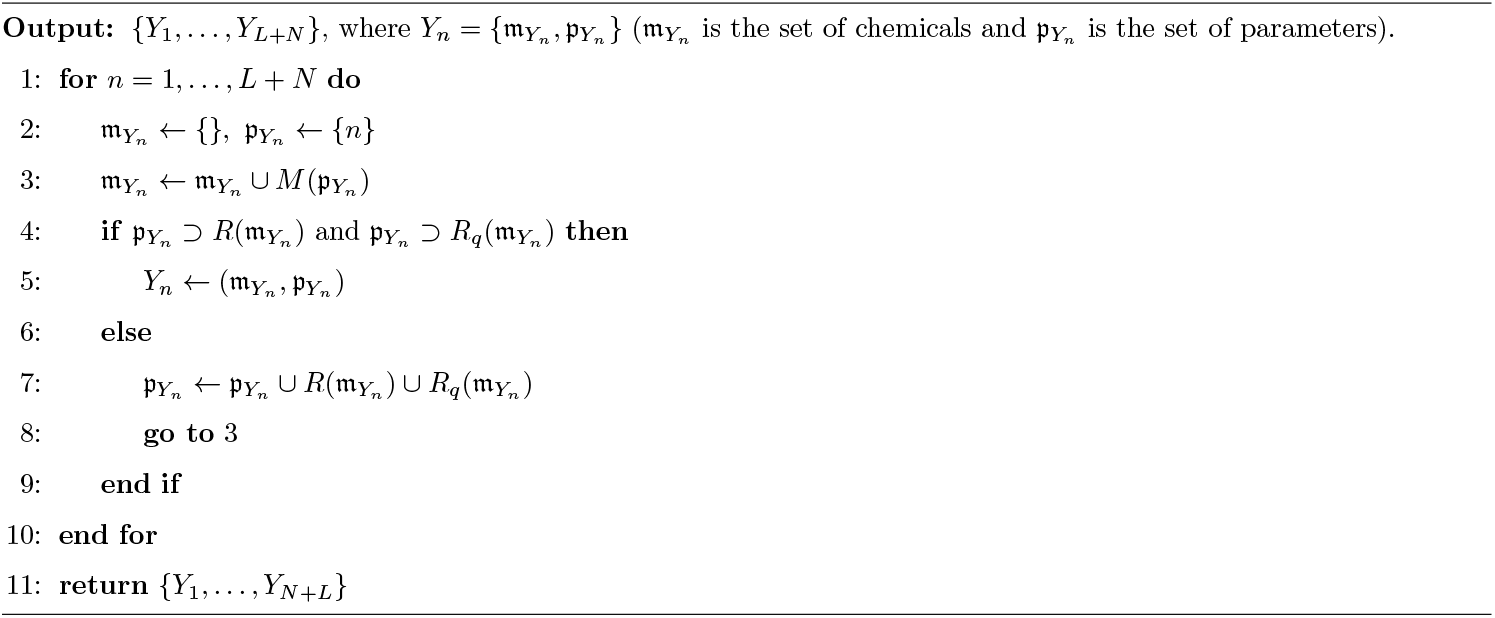

#### Appendix A.4.2 Choice of the cokernel basis

Notably, to find all buffering structures, we have to find response blocks for all possible choices of the basis for ker ***ν***^⊤^, which is not feasible since there are infinitely many ways to select a basis. However, in most cases, finding response blocks under a single basis is sufficient (Proposition S2).

##### Proposition S2.

We consider a regular CRN. Suppose that a basis 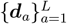 for ker ***ν***^⊤^ can be chosen such that

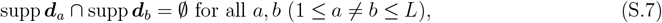

i.e., no two conserved quantities share constituent chemicals. Then, finding response blocks under 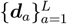 is sufficient to identify all buffering structures.

#### Proof of Proposition S2

Suppose on the contrary that there exists a buffering structure Γ = (𝔪_Γ_, 𝔯_Γ_), which is not a response block under the basis 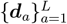 . If supp ***d***_*a*_ ∩ 𝔪_Γ_ = ∅ for all *a* = 1, …, *L*, Γ is a response block regardless of the choice of basis. Therefore, we assume, without loss of generality, that

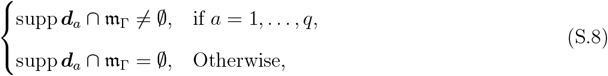

where *q* is the size of the set *{****d***_*a*_ | supp ***d***_*a*_ ∩ 𝔪_Γ_ ≠ ∅*}*. Let 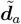 be the projection of ***d***_*a*_ into 𝔪_Γ_. Based on (S.7), 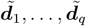 are linearly independent. We denote the basis of ker ***ν*** by 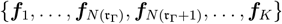, as in Lemma 2. As demonstrated in the proof of Theorem 3, by arranging the orders of the column and row indices of ***A***, we can rewrite ***A*** into the block form as shown in Fig. 6 (Left).

**Fig. 6:**
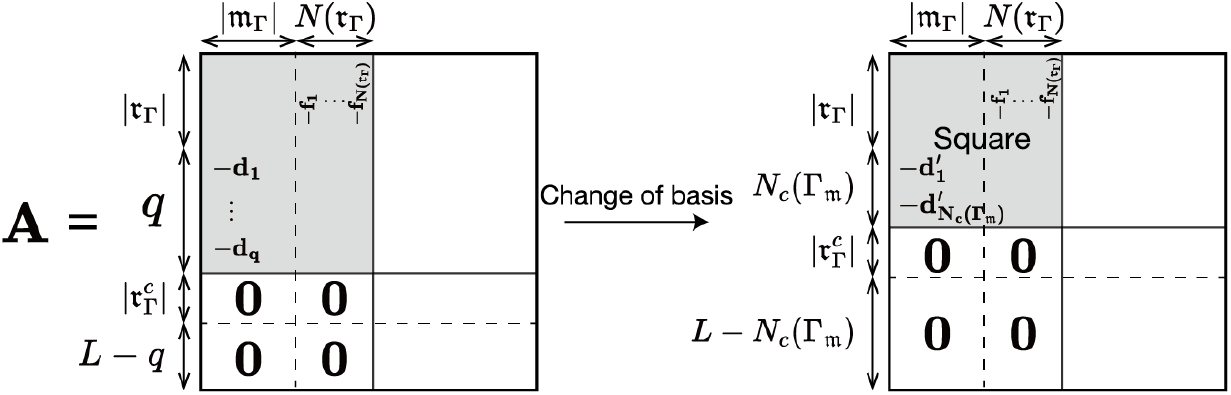
Schematic depiction of the matrix *A* related to the proof of Proposition S2. (Left) By collecting the indices associated with Γ into the upper-left corner, the matrix ***A*** can be a block matrix in which the lower right is the zero matrix. (Right) After the change of basis from 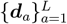 to 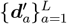, the matrix ***A*** will be a block diagonal matrix.

The structure of block matrices in Fig. 6 (Left) can be obtained by collecting the indices associated with Γ into the upper-left corner: The column indices at the upper left block consist of the chemicals in 𝔪_Γ_ followed by 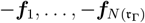, which represent the basis vectors of 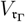 . The row indices consist of the reactions in 𝔯_Γ_ and −***d***_1_, …, −***d***_*q*_. The left block vanishes, similar to the proof of Theorem 3. Regularity of the system indicates |𝔪_Γ_| + *N* (𝔯_Γ_) ≤ |𝔯_Γ_| + *q*. If |𝔪_Γ_| + *N* (𝔯_Γ_) = |𝔯_Γ_| + *q*, the sensitivity matrix ***S*** = −***A***^−1^ is a block-diagonal matrix, which means that Γ is a response block under the basis 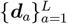. Becasue Γ is not a response block under 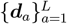, we have

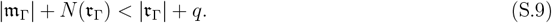

There exsits another basis for ker ***ν***^⊤^, denoted by 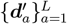, such that 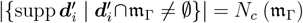, as stated in Theorem S2. Since Γ is a buffering structure,

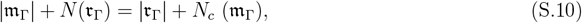

and ***A*** becomes a block diagonal matrix (Fig. 6 (Right)). From (S.9) and (S.10), we have *N*_*c*_ (𝔪_Γ_) *< q*. This means that a change of the basis from 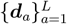 to 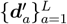 results in a reduction in the number of vectors that have at least one nonzero entry in 𝔪_Γ_. Consequently, there exists *i* (1 ≤ *i* ≤ *L*) such that 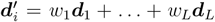 (at least one of *w*_1_, …, *w*_*L*_ is nonzero) and that supp 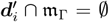. By projecting 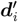 into 𝔪_Γ_, we obtain 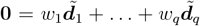 . This contradicts the fact that 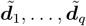 are linearly independent, thereby completing the proof. □

In most CRNs, it is possible to choose the basis for ker ***ν***^*T*^ in such a way that (S.7) holds. In this case, finding response blocks under the single choice of basis is sufficient to find all buffering structures.

##### Example S1.

We consider a pathway, shown in Fig. 7A. We choose the basis for ker ***ν***^*T*^ as ***d***_1_ = (1, 1, 0, 0, 0, 0)^⊤^, ***d***_2_ = (0, 0, 1, 1, 0, 0)^⊤^, ***d***_3_ = (0, 0, 0, 0, 1, 1)^⊤^, which satisfies (S.7). The matrix **A** is

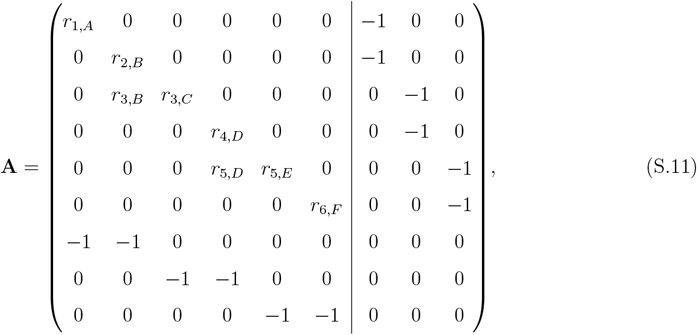

and the sensitivities are determined as

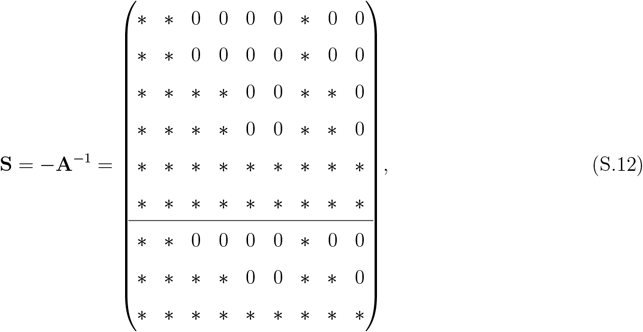

where * represents a nonzero response.

**Fig. 7:**
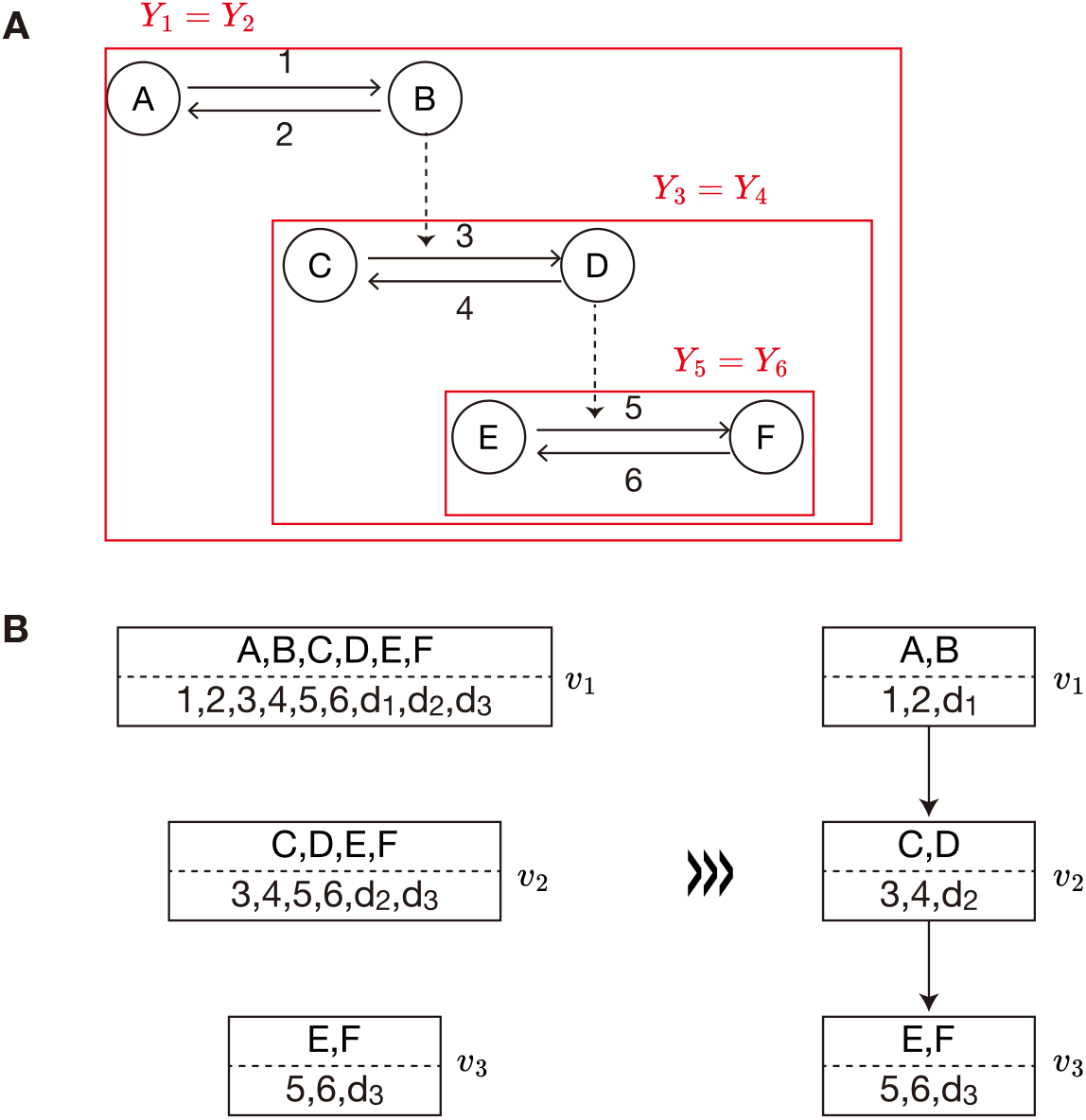
Analysis for a hypothetical network with conserved quantities (Example 1). **A**, Graphical representation of a CRN comprising six chemicals (A,B,C,D,E,F) and six reactions (1,2,3,4,5,6) with three conserved quantities (*d*_1_ : *x*_*A*_ + *x*_*B*_, *d*_2_ : *x*_*C*_ + *x*_*D*_, and *d*_3_ : *x*_*E*_ + *x*_*F*_). Solid lines indicate chemical reactions, while the dashed line indicates active regulation. Each subnetwork enclosed by a red box (*Y*_*i*_) is a buffering structure containing the reaction *i*. **B**, The construction of the hierarchy graph. First, we construct a graph such that each *Y*_*i*_ is assigned to each node *v*_*i*_. Then, we construct the hierarchy graph such that modulating the enzyme activity of reactions within a square box leads to nonzero responses in the chemicals within that box and those in the lower boxes, leaving the other chemicals unaffected.

We obtain

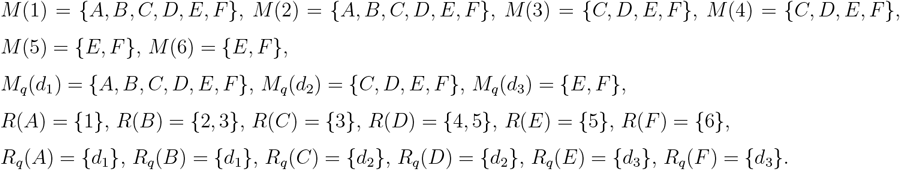

The minimum set of buffering structures are given by the Algorithm 2.

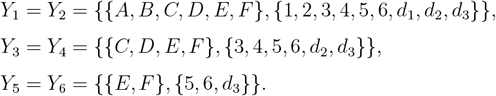

The construction of the hierarchy graph can be done in much the same way as the method described in the main text (Fig. 7B).

##### Remark S2. (Strategy for finding all buffering structures when there is no basis satisfying (S.7))

In some CRNs, choosing a basis satisfying (S.7) is impossible. Even in such cases, we may be able to find all buffering structures using the following strategy.

First, under the specific choice of basis 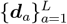 calculated from RREF, we find response blocks Ψ_1_, …, Ψ_*s*_, which are buffering structures from Theorem S3. However, there can be another buffering structure Γ, because there can exist another choice of cokernel basis such that Γ is a response block under the basis. To check this possibility, we examine whether each 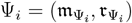 includes a smaller buffering structure (Because the intersection of two buffering structures is also a buffering structure, it is sufficient to check this for each Ψ_*i*_). From the law of localization (Theorem 2), if Ψ_*i*_ includes a hidden buffering structure Γ, perturbations to reaction rate parameters in Γ do not affect the steady-state concentration of chemicals in Ψ_*i*_ *\* Γ. In other words, if some reaction 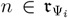 does not affect some chemical 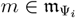, and the nonzero response is not explained by the identified buffering structure Ψ_1_, …, Ψ_*s*_, there is a possibility that Ψ_*i*_ is decomposed into smaller buffering structures. In such cases, changing the basis among 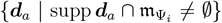 is recommended as it will lead to find the hidden buffering structures.

